# A robust statistical framework for reconstructing genomes from metagenomic data

**DOI:** 10.1101/011460

**Authors:** Dongwan D. Kang, Jeff Froula, Rob Egan, Zhong Wang

## Abstract

We present software that reconstructs genomes from shotgun metagenomic sequences using a reference-independent approach. This method permits the identification of OTUs in large complex communities where many species are unknown. Binning reduces the complexity of a metagenomic dataset enabling many downstream analyses previously unavailable. In this study we developed MetaBAT, a robust statistical framework that integrates probabilistic distances of genome abundance with sequence composition for automatic binning. Applying MetaBAT to a human gut microbiome dataset identified 173 highly specific genomes bins including many representing previously unidentified species.

## INTRODUCTION

High throughput metagenome shotgun sequencing is a powerful tool to study microbial communities directly taken from their environment, thereby avoiding the requirement for cultivation or the biases that may arise from it. Most of the microbial diversity has not been isolated due to the technical difficulties in doing so; thus, most of the microbiome world is unknown. Detecting individual draft genomes in a process called metagenome binning (1,2) provides a substitute for direct isolation. Generally, binning efficiency is dependent on the length of the metagenomic sequences being binned; therefore, the first step may be the assembly of the reads, typically by a *de bruijn* graph based short read assembler (3,4). Even draft genomes lacking full contiguity can approximate full genomes in many aspects as they can contain a near full set of genes.

The large number of metagenomic studies and the sheer volume of sequence being produced from each gives binning tools the crucial role of reducing data complexity opening up the possibility for more downstream analyses. Binning is superior to non-binning methods of annotating metagenomes, like rRNA and marker gene identification, since it is a more comprehensive, automatable, scalable and simple approach. Assuming the accuracy is on par with more manual protocols, automated binning software will also make metagenome annotation cheaper. Considering that a single sample of soil may contain 18,000 unique organisms (5) and that many metagenomic studies require comparing temporal or spatial communities, it is easy to see the advantages of automated binning software.

The large number of assembled contigs from a complex community creates a unique challenge for metagenome binning. Two classes of binning approaches have been developed to tackle this challenge (reviewed in (1)). Supervised binning approach takes advantage of known genomes and relies on either sequence homology or a composition signature for binning (6,7). This approach does not work well on environmental samples where many of the microbes in these communities do not have close relatives with known genomes. In contrast, the unsupervised approach relies on either discriminative sequence composition (8,9) or species (or genomic fragments) abundance (10–14) or both (15–17). However, genomes obtained from complex communities using this approach contain many contaminants (2), presumably due to lack of robust statistical measures.

Our software, MetaBAT (<u>Metag</u>enome Binning with <u>A</u>bundance and <u>T</u>etra-nucleotide frequencies) not only combines tetra-nucleotide frequency (TNF) analysis with abundance analysis but leverages the variation in species abundance between samples. So if two genomes have the same abundance in one sample but significantly different abundances in another sample, they have the chance to be separated into two bins. Each additional sample will reinforce this separation. Genome abundance variation can be due to biologically explained differences or just an artifact of the sample collection itself. The power of this technique depends on the relative inter-sample variation versus intra-sample variation of a genome.

## MATERIAL AND METHODS

### An overview of MetaBAT software and its probabilistic models

Briefly, MetaBAT does the following. For each pair of contigs in a metagenome assembly, it first calculates their probabilistic distance: 1) based on the tetranucleotide frequency and 2) based on abundance. The two distances are integrated into one composite distance. All the pairwise distances form a matrix, which then is subject to a modified k-medoid clustering algorithm to iteratively bin the contigs into single genomes (Figure 1).

**Figure 1.**
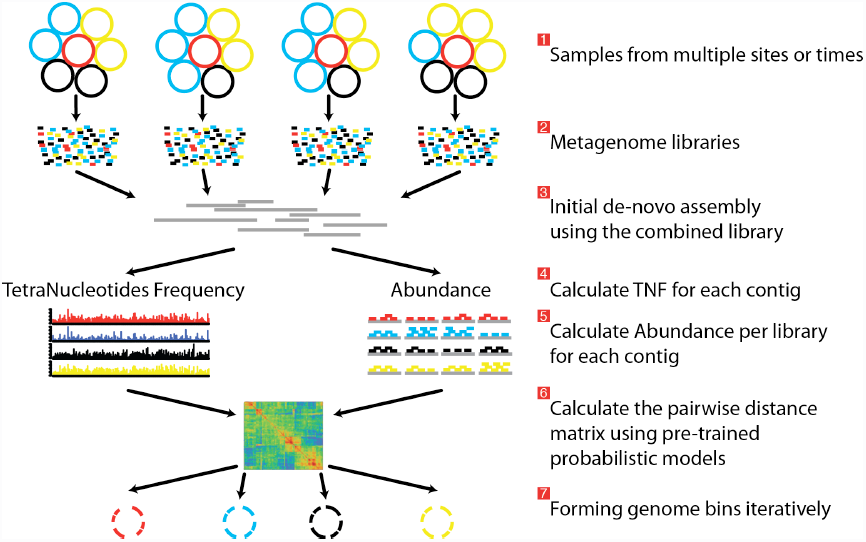
Overview of the MetaBAT pipeline. Three preprocessing steps before MetaBAT is applied: 1) A typical metagenome experiment may contain many spatial or time-series samples, each consisting of many different genomes (different color circles). 2) Each sample is sequenced by next-generation sequencing technology to form a sequencing library with many short reads. 3) The libraries may be combined before de novo assembly. After assembly, the reads from each sample must be aligned in separate BAM files. MetaBAT then automatically performs the remaining steps: 4) For each contig pair, a tetranucleotide frequency distance probability (TDP) is calculated from a distribution modelled from 1,414 reference genomes. 5) For each contig pair, an abundance distance probability (ADP) across all the samples is calculated. 6) The TDP and ADP of each contig pair are then combined, and the resulting distance for all pairs form a distance matrix. 7) Each bin will be formed iteratively and exhaustively from the distance matrix. See below for more details.

We use tetranucleotide frequency as sequence composition signatures, as it has been previously shown that different microbial genomes have distinct TNF biases (18–21). To empirically derive a distance to discriminate TNFs of different genomes (TDP), we calculated the likelihood of inter- and intra-species Euclidean distance by using 1,414 unique, complete genome references from NCBI (Figure 2a). To evaluate the effect of contig sizes on inter-species distance, we obtained posterior probability distributions of inter-species distance with several fixed sizes and observed better inter-species separation as contig size increases (Figure 2b). As contigs in real metagenome assemblies have various sizes, we then modeled TDP between contigs of different sizes by fitting a logistic function to reflect the dynamic nature of non-linear relationship between Euclidean TNF distance and TDP in different contig sizes. The results (Figure 2, c and d) suggested that the values of two parameters of the model, b and c, are very unstable if the size of either contig is very small (< 2kb) and it should be cautious to allow smaller contigs included in the binning.

**Figure 2.**
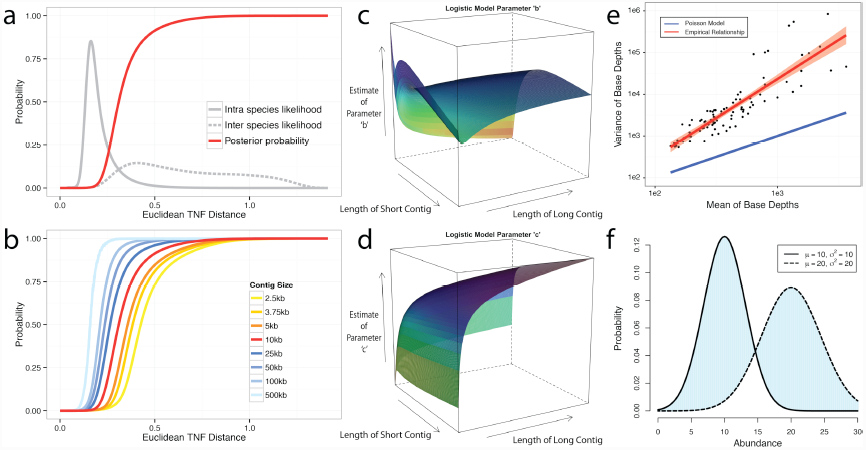
Probabilistic modeling of TNF and Abundance distances. **a-d**) TNF distance modeling. **a**) Empirical probabilities of intra- (solid gray line) or inter- (dotted gray line) species Euclidean TNF distance are estimated from sequenced genomes. The posterior probability of two contigs originated from different genomes given a TNF distance is shown as a red solid line. All probabilities are calculated using a fixed contig size of 10kb. **b**) Different posterior inter-species probabilities for two equal-size contigs under various contig sizes. **c, d**) The estimation of parameters for a logistic curve with two contigs of different sizes. × and y axis represent the lengths of short and long contig, respectively, and z axis represents the estimates of each parameter *b* or *c* in a logistic curve, TDP = 1/(1+exp(-(*b*+*c*\*TNF))), where TNF and TDP represents the Euclidean TNF distance and probabilistic TNF distance, respectively. **e-f**) Abundance distance modeling. **e**) The relationship between mean and variance of base depths (coverage) which were shown in x and y axis, respectively. Each dot represents this relationship in each genome, which calculated by median of mean and variance of the coverage. Theoretical Poisson model was shown as blue line and normal model was shown as red line. **f**) Probabilistic abundance distance between two contigs. The shaded area represents the abundance distance between two contigs in a given library.

Although contigs originating from the same genome are expected to have similar sequence coverage, i.e. genome abundance, the coverage of contigs can vary significantly, within a library, due to biases originated from the current sequencing technology (22–25). As illustrated in Figure 2e, the observed coverage variance derived from data consisting of isolate genome sequencing projects (total 99 from IMG Database (26), henceforth referred as the IMG dataset) significantly deviate from the theoretical Poisson distribution, consistent with the previous notion that both the variance and the mean should be modeled (27). For computational convenience, we chose the normal distribution as an approximation since it fits the observation much better (Figure 2e). To compute the abundance distance of two contigs in one sample, we use the area not shared by their inferred normal distributions with given coverage mean and variance (Figure 2f). A geometric mean of the distances for all samples is used for the final abundance distance probability (ADP) of two contigs. In addition, we applied a progressive weight mechanism to adjust the relative strength of the information from abundance distance, meaning that we put more weight on abundance distance when it was calculated from many samples (see below).

We then integrate TDP and ADP of each contig pairs as the following:

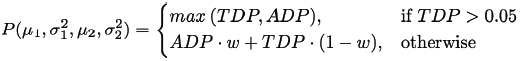

, where w = min [ log(n+1) / log(m+1), α]. n, m, and α represent the number of samples, a large number (100 as the default), and the maximum weight of ADP (0.9 as the default), respectively. For instance, in the default setting, the weight would be about 0.5 when the number of samples is 10 and TDP is less than 0.05. The resulting distance matrix is used for binning (see below).

### Tetranucleotide Frequency Probability Distance (TDP)

To establish empirical probabilities of intra- and inter-species for tetranucleotide frequency (TNF) distance, we compiled a set of unique, complete genomes from NCBI database, and shredded the genomes into fragments ranging from 2.5kb to 500kb and obtained 1 billion random contig pairs from within or between genomes using 1,414 unique complete genomes in NCBI. The empirical posterior probability that two contigs are from different genomes is given as the following:

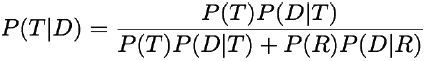

, where T or R represent cases where two contigs are from different (inter) or the same (intra) species, respectively. D is the Euclidean TNF distance between two contigs. The same uninformative priors of T and R were chosen. In reality, P(T) is expected to be much bigger than P(R), thus we set P(T) = 10* P(R) as the default implementation to adjust the possible under-sampling issue in inter species distance.

The TDP of contig pairs with different sizes is approximated using logistic regression:

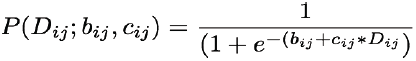

, where *D*_*ij*_ represents a Euclidean TNF distance between contig *i* and *j*. b and c, the two parameters for the logistic regression, are estimated from the empirical data.

### Abundance Distance Probability (ADP)

In complex microbial communities with many related species, homologous reads from one species can map to another and this leads to inaccurate contig abundance measurement. We therefore adopted a very stringent criterion by requiring mapped reads to have at least 97% identity over their entire length. To avoid the poor mapping at the contig edges, we also exclude one read length from each end of a contig for abundance calculation. MetaBAT accepts sorted BAM files as the default input (28).

The probabilistic abundance distance was calculated as follows: Suppose two contigs have the mean coverage of µ_1_ and µ_2_ and the variances of σ_1_^2^ and σ_2_^2^, then we defined the abundance distance as the non-shared area of two normal distribution of *Ν*(µ_1_, σ_1_^2^) and *Ν*(µ_2_, σ_2_^2^):

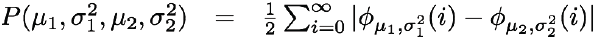

, where ϕ represents a normal distribution having two parameters µ and σ^2^. Numerically this can be simplified using cumulative distribution functions as follows assuming σ_2_^2^ is greater than σ_1_^2^:

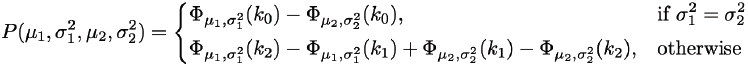

, where Φ represents a cumulative normal distribution, and

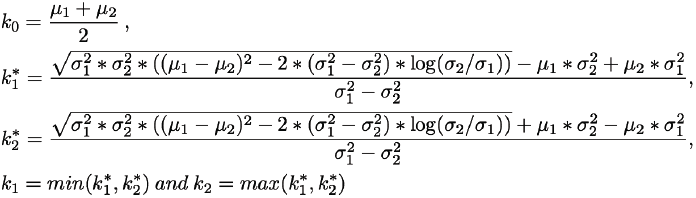

To combine multiple abundance probabilities across different samples, we calculated the geometric mean of probabilities:

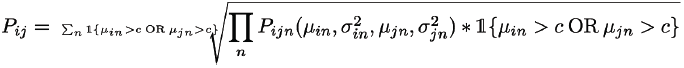

, where P_ijn_ represents the probability calculated from two abundances µ_in_ and µ_jn_, and c represents a cutoff for reasonable minimum abundance for a contig.

As metagenome assemblies contain many small contigs, whether or not to include them is a dilemma: including small contigs will likely improve the genome completeness, at a cost in genome quality because their larger abundance variations make it harder to bin them correctly. We tried to empirically determine a reasonable contig size cutoff by plotting the ratio of mean and variance from the IMG single genome dataset. Although for most of genomes variances are much larger than means, their ratio becomes stabilized after contig size increases to 2.5kb (Supplementary Figure S1).

### Iterative Binning

We modified the k-medoid clustering algorithm (29) to eliminate the requirement of the user input for *k* and to reduce search space for efficient binning. Specifically, the binning algorithm works as the following:

1. Find a seed contig (e.g. having the largest coverage), and set it as the initial medoid.
2. Recruit all other contigs within a cutoff distance (i.e. sensitivity parameters p1 and p2) to the seed.
3. Find a new medoid out of all member contigs.
4. Repeat 1-3 until there are no further updates to the medoid. These contigs form a bin.
5. For the rest of the contigs, repeat 1-4 to form more bins until no contigs are left.
6. Keep large bins (e.g. >200kb), and dissolve all other bins into free contigs.
7. For each of the large bins, recruit additional members from free contigs that are within another cutoff distance (i.e. sensitivity parameter p3) to any of the member contigs, to form the final genome bin.

## RESULTS

### Construction of a synthetic metagenomic assembly dataset for benchmarking MetaBAT

A metagenome dataset from the MetaHIT consortium (henceforth referred to as the MetaHIT dataset) (30) was chosen to benchmark MetaBAT because it contained a large number of samples and the community contains many species with reference genomes. We used the largest 256 metagenomic samples from a total of 262 MetaHIT samples. To find a reference genome set, we downloaded 9,673 reference bacterial genomes (complete and draft) from NCBI and mapped the MetaHIT metagenome reads to them using BBMAP (http://bbmap.sourceforge.net) with default parameters. 240 genomes can be found in one or more samples (Supplementary Table S2). Only large and relatively abundant genomes (>200kb, total coverage > 5X in all samples combined and > 2X in at least 1 sample) are selected to form the reference genome set. These reference genomes, 237 in total, were then shredded into contigs of random sizes (> 2.5 kb) to mimic real metagenome assemblies without assembly errors or biases. In each of the 10 simulations, on average we generated 96,201 contigs. These “error-free” metagenome contigs, along with their parental reference genomes representing true answers, were used in the following analyses to benchmark MetaBAT’s binning performance.

We used three criteria to measure binning performance. The “completeness” criterion measures the ability to bin all contigs from the same genome; the “precision” criterion measures the binning specificity, or avoiding contigs from other “contaminant” genomes; and finally, “community completeness” criterion measures the ability to recover all the genomes that are present in the community. The specific definition of these criteria can be found in the supplemental materials.

### Binning performance on error-free metagenomic assemblies

The above “error-free” metagenome contigs derived from 231 reference genomes and the 256 experimental metagenome sequence datasets were used to benchmark MetaBAT. We first tested MetaBAT’s efficiency with varying numbers of samples. Although the sample number varied, the number of sequences was kept constant. Overall the binning performance increased as the number of samples increased (Figure 3a). The binned single genomes rarely contain “contaminant” contigs even with a small number of samples, although the genomes become more complete when more samples were used. This robust high precision feature is desirable since most current metagenome experiments have only a few samples. In addition, more near complete genomes are binned (community completeness metric) as the number of sample increases (Supplementary Figure S2).

**Figure 3.**
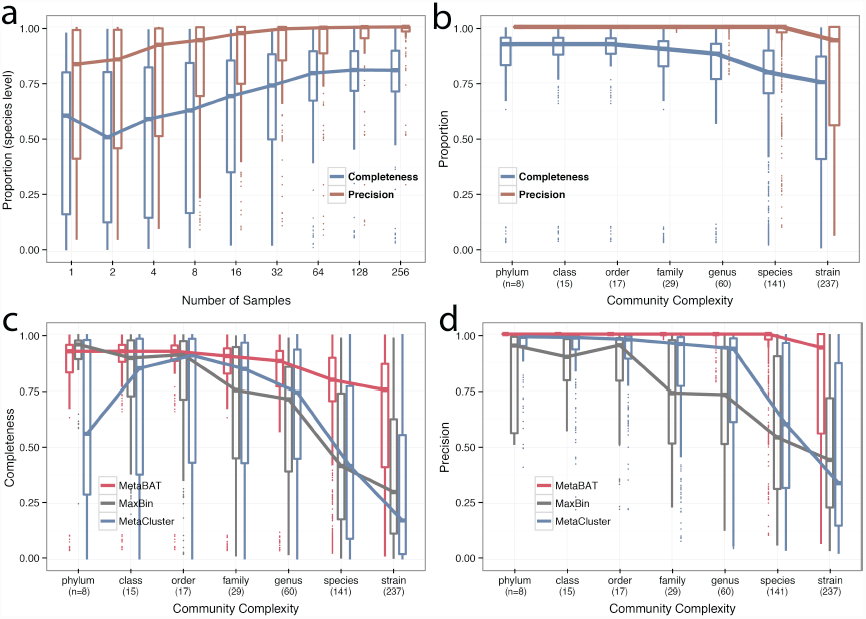
Binning performance on synthetic metagenomic assemblies. **a**) Boxplots of the completeness metric (purple) and precision metric (red) of MetaBAT on different number of samples. **b**) The completeness and precision performance of MetaBAT at different levels of community complexity. **c,d**) Binning performance comparison between MetaBAT, MaxBin and MetaCluster for completeness (**c**) and precision (**d**) on the same dataset.

Next we varied the community complexity and tested MetaBAT’s performance. The reference genomes in the MetaHIT dataset belong to 8 phyla, 15 classes, 17 orders, 29 families, 60 genus, 141 species, and 237 strains. We constituted a synthetic community with the lowest complexity by choosing one genome from each phylum. Similarly, six more communities with increasing complexity were synthesized by choosing one genome from each class, order, family, genus, species or strain level. The community including all 237 strains is the most complex (Supplementary Table S2). When the number of samples was fixed to 256 for all synthetic communities, MetaBAT displays robust performance as community complexity increases (Figure 3b), predicting near complete genomes without significant contaminants at species level or above (100% median precision). In addition, 70% of the species from the community are recovered (Supplement Figure S2). Remarkably, MetaBAT can discriminate among different strains within the same species with 94% precision, presumably due to its implementation of robust statistics metrics.

Using the same synthetic metagenome assemblies we compared the performance of MetaBAT against two other automatic binning methods using their default settings, MaxBin (31) and MetaCluster (32). MetaBAT outperforms both methods in completeness, precision and community completeness when the communities are complex (species level mixing or above, Figure 1c,d and Supplement Figure S2).

### Binning performance on real world metagenome assemblies

To validate the performance of MetaBAT on real metagenomic data sets, we first combined all the sequences from the 262 samples in the MetaHIT dataset and assembled them using Ray Meta Assembler (33). We selected 60,619 contigs longer than 2.5kb for binning. To systematically explore the trade-offs between precision and completeness, we ran MetaBAT under 4 pre-specified parameter sets: very sensitive, sensitive, specific, and very specific.

As shown in Figure 4a, with the default ‘sensitive’ setting, among the 65 bins that have matching reference genomes, MetaBAT forms 37 near complete genomes with a mean size of 2.2Mb and 26 partial genomes having mean size of 1.0Mb. For the two bins that have poor reference genome matches (∼20%), we could not distinguish whether they represented chimeric bins, or a distant relative of the matching reference. We found larger bins under the ‘very sensitive’ setting at the cost of more contamination, whereas the ‘very specific’ setting had no contamination but smaller bins with fewer complete genomes.

**Figure 4.**
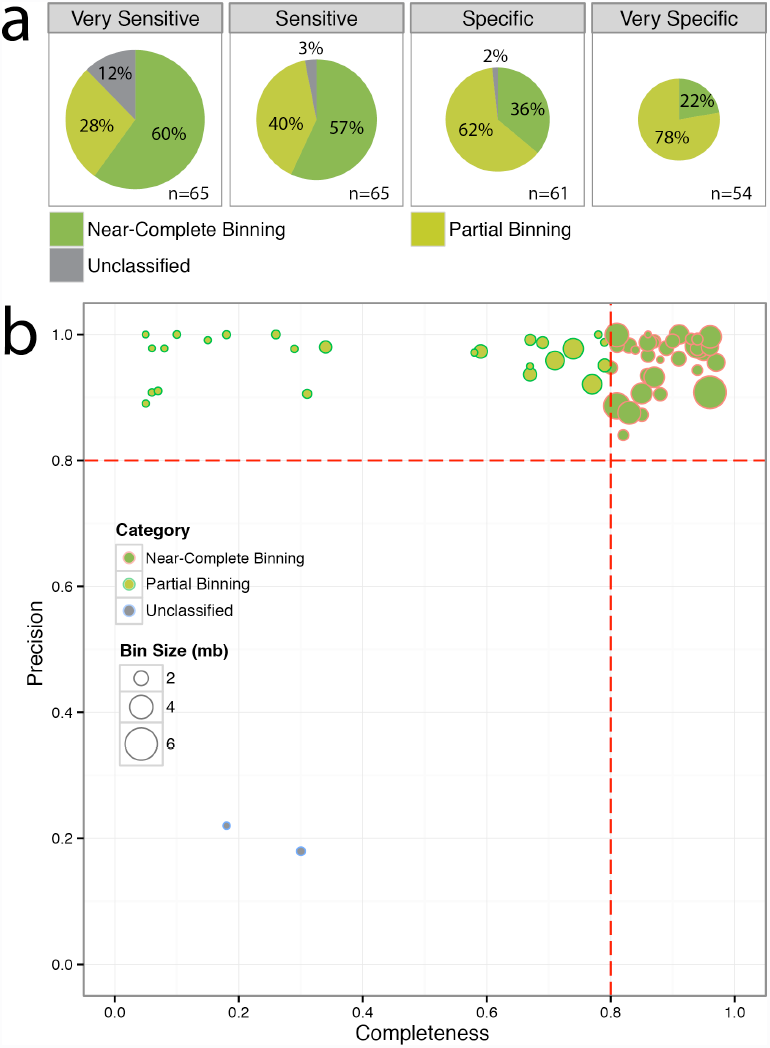
Binning results on the MetaHIT dataset. **a**) Performance comparison of 4 pre-specified parameter settings in MetaBAT, estimated by genome bins with known reference. Near-Complete Genomes: completeness >= 80% and precision >= 80%; Partial Genomes: precision >= 80% and completeness < 80%; Unclassified: 10% < precision < 80%. The size of each pie chart corresponds to the total bases of all the genome bins. *n*: number of bins with reference. **b**) Distribution of genomes binned under the default ‘Sensitive’ setting.

Further phylogenic analyses of the 108 putative genomes under the ‘sensitive’ setting suggest they encompass various taxonomic levels, including 16 new species only identifiable at the family level (Supplementary Table S1).

## DISCUSSION

In conclusion, we developed robust metagenome binning software, MetaBAT, and demonstrated its superior performance to reconstruct genomes from both synthetic and real world metagenome datasets. Applying MetaBAT to a complex human microbiome recovers hundreds of high quality genomes including many putative new species.

Although we showed that MetaBAT achieves robust binning on both synthetic and real metagenome assemblies, there are several considerations that users should keep in mind before applying MetaBAT to their metagenome assemblies. First, what would be the minimum number of samples for a successful binning? It is not necessary to have multiple samples to run the software. MetaBAT can run with one sample and even accommodate TDP-only binning without abundance information. However, as shown in Figure3a, our general advice is that more samples would achieve better binning results. The number of samples in current metagenome studies is often decided by scientific aims other than best binning results. Even if we were to design an experiment for best binning results, the minimum number of samples would also depend on the degree of community structural differences among the samples.

Our general expectation is that, the greater the abundance of a target species varies among samples, the more likely MetaBAT will produce a good genome bin for this species. Second, we do not expect binning to work well with poor metagenome assemblies, e.g. assemblies including many small contigs less than 2.5kb. The distance metrics computed for small contigs will not be very reliable. Finally, the metaHIT dataset is unique as it contains many species with known reference genomes. In many environmental metagenome samples it may not be easy to evaluating the quality of the resulting genome bins, as most of their member species are unknown.

Since we are using an unsupervised approach for binning without any optimization towards any specific species, the binning performance estimated from known reference genomes should provide an unbiased estimation about the quality of the novel ones. Previously *de novo* metrics have been introduced to evaluate the quality of genome bins without reference matches (34–36). However, based on our own experience, the marker gene based (core genes and single copy genes) quality assessment is not very accurate. For example, clustered core genes may cause overestimation of the completeness, and the fact that some bacteria may lack significant number of core genes will lead to underestimation of completeness. Moreover, we observed multiple alleles of a single gene were assembled into separate contigs, which leads to overestimation of the contamination.

Implemented in C++, MetaBAT runs very fast. It took only 1 hour to process the MetaHIT dataset (60,619 contigs and 262 bam files) with 16 CPU cores and 2.7 GB memory. The adoption of multiple parameter settings also makes MetaBAT versatile for different analysis requirements for sensitivity and specificity. The software is fully automated and easy to use. It is open source and available to download (https://bitbucket.org/berkeleylab/metabat).

## ACKNOWLEDGEMENT

The authors thank Drs. Matt Blow, Rex Malmstrom and Tanja Woyke for their stimulating discussions and critical comments.

## REFERENCES

1. Mande S.S., Mohammed, M.H. and Ghosh, T.S. (2012) Classification of metagenomic sequences: methods and challenges. Briefings in bioinformatics, 13, 669–681.

2. Mavromatis, K., Ivanova, N., Barry, K., Shapiro, H., Goltsman, E., McHardy, A.C., Rigoutsos, I., Salamov, A., Korzeniewski, F., Land, M. et al. (2007) Use of simulated data sets to evaluate the fidelity of metagenomic processing methods. Nature methods, 4, 495–500.

3. Pevzner, P.A. and Tang, H. (2001) Fragment assembly with double-barreled data. Bioinformatics, 17 Suppl 1, S225–233.

4. Pevzner, P.A., Tang, H. and Waterman, M.S. (2001) An Eulerian path approach to DNA fragment assembly. Proc. Natl. Acad. Sci. USA, 98, 9748–9753.

5. Daniel, R. (2005) The metagenomics of soil. Nature reviews. Microbiology, 3, 470–478.

6. Krause, L., Diaz, N.N., Goesmann, A., Kelley, S., Nattkemper, T.W., Rohwer, F., Edwards, R.A. and Stoye, J. (2008) Phylogenetic classification of short environmental DNA fragments. Nucleic acids research, 36, 2230–2239.

7. Wu, M. and Eisen, J.A. (2008) A simple, fast, and accurate method of phylogenomic inference. Genome biology, 9, R151.

8. Teeling, H., Waldmann, J., Lombardot, T., Bauer, M. and Glockner, F.O. (2004) TETRA: a web-service and a stand-alone program for the analysis and comparison of tetranucleotide usage patterns in DNA sequences. BMC bioinformatics, 5, 163.

9. Yang, B., Peng, Y., Leung, H.C., Yiu, S.M., Chen, J.C. and Chin, F.Y. (2010) Unsupervised binning of environmental genomic fragments based on an error robust selection of l-mers. BMC bioinformatics, 11 Suppl 2, S5.

10. Wu, Y.W. and Ye, Y. (2011) A novel abundance-based algorithm for binning metagenomic sequences using l-tuples. J. Comp. Biol., 18, 523–534.

11. Cotillard, A., Kennedy, S.P., Kong, L.C., Prifti, E., Pons, N., Le Chatelier, E., Almeida, M., Quinquis, B., Levenez, F., Galleron, N. et al. (2013) Dietary intervention impact on gut microbial gene richness. Nature, 500, 585–588.

12. Qin, J., Li, Y., Cai, Z., Li, S., Zhu, J., Zhang, F., Liang, S., Zhang, W., Guan, Y., Shen, D. et al. (2012) A metagenome-wide association study of gut microbiota in type 2 diabetes. Nature, 490, 55–60.

13. Nielsen H.B., Almeida, M., Juncker, A.S., Rasmussen, S., Li, J., Sunagawa, S., Plichta, D.R., Gautier, L., Pedersen, A.G., Le Chatelier, E. et al. (2014) Identification and assembly of genomes and genetic elements in complex metagenomic samples without using reference genomes. Nature biotechnology.

14. Le Chatelier, E., Nielsen, T., Qin, J., Prifti, E., Hildebrand, F., Falony, G., Almeida, M., Arumugam, M., Batto, J.M., Kennedy, S. et al. (2013) Richness of human gut microbiome correlates with metabolic markers. Nature, 500, 541–546.

15. Albertsen, M., Hugenholtz, P., Skarshewski, A., Nielsen, K.L., Tyson, G.W. and Nielsen, P.H. (2013) Genome sequences of rare, uncultured bacteria obtained by differential coverage binning of multiple metagenomes. Nature biotechnology, 31, 533–538.

16. Sharon, I., Morowitz, M.J., Thomas, B.C., Costello, E.K., Relman, D.A. and Banfield, J.F. (2013) Time series community genomics analysis reveals rapid shifts in bacterial species, strains, and phage during infant gut colonization. Genome research, 23, 111–120.

17. Wrighton K.C., Thomas, B.C., Sharon, I., Miller, C.S., Castelle, C.J., VerBerkmoes N.C., Wilkins, M.J., Hettich, R.L., Lipton, M.S., Williams, K.H. et al. (2012) Fermentation, hydrogen, and sulfur metabolism in multiple uncultivated bacterial phyla. Science, 337, 1661–1665.

18. Pride D.T., Meinersmann, R.J., Wassenaar, T.M. and Blaser, M.J. (2003) Evolutionary implications of microbial genome tetranucleotide frequency biases. Genome research, 13, 145–158.

19. Teeling, H., Meyerdierks, A., Bauer, M., Amann, R. and Glockner, F.O. (2004) Application of tetranucleotide frequencies for the assignment of genomic fragments. Environmental microbiology, 6, 938–947.

20. Mrazek, J. (2009) Phylogenetic Signals in DNA Composition: Limitations and Prospects. Mol Biol Evol, 26, 1163–1169.

21. Saeed, I., Tang, S.L. and Halgamuge, S.K. (2012) Unsupervised discovery of microbial population structure within metagenomes using nucleotide base composition. Nucleic acids research, 40, e34.

22. Harismendy, O., Ng, P.C., Strausberg, R.L., Wang, X., Stockwell, T.B., Beeson, K.Y., Schork, N.J., Murray, S.S., Topol, E.J., Levy, S. et al. (2009) Evaluation of next generation sequencing platforms for population targeted sequencing studies. Genome biology, 10, R32.

23. Nakamura, K., Oshima, T., Morimoto, T., Ikeda, S., Yoshikawa, H., Shiwa, Y., Ishikawa, S., Linak, M.C., Hirai, A., Takahashi, H. et al. (2011) Sequence-specific error profile of Illumina sequencers. Nucleic acids research, 39, e90.

24. Benjamini, Y. and Speed, T.P. (2012) Summarizing and correcting the GC content bias in high-throughput sequencing. Nucleic acids research, 40, e72.

25. Ross M.G., Russ, C., Costello, M., Hollinger, A., Lennon, N.J., Hegarty, R., Nusbaum, C. and Jaffe, D.B. (2013) Characterizing and measuring bias in sequence data. Genome biology, 14, R51.

26. Markowitz, V.M., Chen, I.M., Palaniappan, K., Chu, K., Szeto, E., Grechkin, Y., Ratner, A., Jacob, B., Huang, J., Williams, P. et al. (2012) IMG: the Integrated Microbial Genomes database and comparative analysis system. Nucleic acids research, 40, D115–122.

27. Clark, S.C., Egan, R., Frazier, P.I. and Wang, Z. (2013) ALE: a generic assembly likelihood evaluation framework for assessing the accuracy of genome and metagenome assemblies. Bioinformatics, 29, 435–443.

28. Li, H., Handsaker, B., Wysoker, A., Fennell, T., Ruan, J., Homer, N., Marth, G., Abecasis, G., Durbin, R. and Genome Project Data Processing, S. (2009) The Sequence Alignment/Map format and SAMtools. Bioinformatics, 25, 2078–2079.

29. Kaufman, L. and Rousseeuw, P. (1987) Clustering by means of medoids. North-Holland.

30. Qin, J.J., Li, R.Q., Raes, J., Arumugam, M., Burgdorf, K.S., Manichanh, C., Nielsen, T., Pons, N., Levenez, F., Yamada, T. et al. (2010) A human gut microbial gene catalogue established by metagenomic sequencing. Nature, 464, 59–U70.

31. Wu, Y.-W., Tang, Y.-H., Tringe, S.G., Simmons, B.A. and Singer, S.W. (2014) MaxBin: an automated binning method to recover individual genomes from metagenomes using an expectation-maximization algorithm. Microbiome, 2, 26.

32. Wang, Y., Leung, H.C., Yiu, S.M. and Chin, F.Y. (2012) MetaCluster 5.0: a two-round binning approach for metagenomic data for low-abundance species in a noisy sample. Bioinformatics, 28, i356–i362.

33. Boisvert, S., Raymond, F., Godzaridis, E., Laviolette, F. and Corbeil, J. (2012) Ray Meta: scalable de novo metagenome assembly and profiling. Genome biology, 13, R122.

34. Mende D.R., Sunagawa, S., Zeller, G. and Bork, P. (2013) Accurate and universal delineation of prokaryotic species. Nature methods, 10, 881–884.

35. Rinke, C., Schwientek, P., Sczyrba, A., Ivanova, N.N., Anderson, I.J., Cheng, J.F., Darling, A., Malfatti, S., Swan, B.K., Gies, E.A. et al. (2013) Insights into the phylogeny and coding potential of microbial dark matter. Nature, 499, 431–437.

36. Shi, W., Moon, C.D., Leahy, S.C., Kang, D., Froula, J., Kittelmann, S., Fan, C., Deutsch, S., Gagic, D., Seedorf, H. et al. (2014) Methane yield phenotypes linked to differential gene expression in the sheep rumen microbiome. Genome research.

